# The repercussions of timing in the invasion of synthetic bacterial communities

**DOI:** 10.1101/2024.03.24.586461

**Authors:** Keven D. Dooley, Joy Bergelson

## Abstract

Microbial communities regularly experience ecological invasions that can lead to changes in composition and function. Factors thought to affect the outcome of invasions, like diversity and resource use, vary over the course of community assembly, potentially altering susceptibility to arriving invaders. We used synthetic bacterial communities to evaluate the success and impact of invasions occurring at different times during the community assembly process. Fifteen distinct communities were subjected to each of three bacterial invaders at the initial assembly of the community (“initial”), 24 hours into community assembly (“early”), when the community was still undergoing transient dynamics, and 7 days into community assembly (“late”), once the community had settled into its final composition. Communities were passaged daily and characterized through sequencing after reaching a stable composition. Invasions were most successful and had their largest effect on composition when they occurred before a community had settled into a stable composition. Surprisingly, we found instances where an invader was ultimately excluded yet had profound and long-lasting effects on invaded communities. We also found that common community members were more greatly impacted by invaders than rare community members. Higher invasion success and impact were associated with lower community resource use efficiency, which varied throughout assembly. Our results demonstrate that microbial communities experiencing transient community dynamics are more prone to invasion, a finding relevant to efforts to modify the composition of microbial communities.

## Introduction

Microbial communities are ubiquitous and of great importance to human health (1), agriculture (2), and industry (3). As such, many efforts are underway to design or modify microbial communities that assume a desired composition or perform a desired function. But, like all ecological communities, microbial communities are exposed to environmental fluctuations and migration that can affect community composition and function (4,5). This represents a challenge for our efforts to manipulate microbial communities to serve our own ends.

Take, for example, efforts to design a microbial community that performs a desired function. As has often been observed, communities that perform as designed *in vitro* or in a host under well controlled conditions frequently shift in composition and function when exposed to the complexity of a natural environment (4,6,7). Or consider the engraftment of a probiotic bacterium in the gut microbiome. The potential positive effect of that probiotic on host health is irrelevant if it cannot persist in the host microbiome (8,9). Both these scenarios deal with the complexities of ecological invasions and highlight the importance of understanding when and why communities become vulnerable to invaders.

Invasion ecology concerns the establishment and impact of novel species on ecological communities. There is a well-established relationship between community diversity and invasibility, wherein more diverse communities are often more resistant to invasion (10–14). This relationship has been attributed to two mechanisms, 1) the “sampling effect” (11,15–17), where a more diverse community is more likely to include species that directly compete with an invader, and 2) a positive relationship between community diversity and resource use efficiency (10,15,18), wherein a more diverse community is better able to use the resources available in the environment, leaving little available niche space for an invader to occupy. However, communities passing through transient dynamics can suffer from elevated competition and reduced resource use efficiency (19,20), suggesting that these highly diverse communities might be less resistant to invasion. It is clear that diversity and resource use are dynamic over the course of community assembly as species are lost and change in abundance, and such variation may affect the invasibility of assembling communities.

Here, we seek to experimentally disentangle the roles of community richness, resource use, and the assembly process in determining resistance to invasion. We posit that the timing of an invasion impacts its outcome due to changes in the diversity (richness) and function (resource use) of communities as they assemble, and reassemble, over time. We consider the “outcome” of an invasion as both the success/failure of the invader to persist in a community over time and the effect of the invader on the composition of the community.

## Materials and Methods

### Bacterial isolates and reference genomes

All bacterial isolates were originally isolated from the leaves of wild or field grown *Arabidopsis thaliana* in the midwestern states of the USA (IL, IN, MI). The isolate names, taxonomic information, and assembly information are presented in Supplementary Table 1. Genomes are published on the NCBI whole genome repository (accessions: PRJNA953780, PRJNA1073231).

### Arabidopsis leaf medium (ALM)

*Arabidopsis thaliana* (KBS-Mac-74, accession 1741) plants were grown in the University of Chicago greenhouse from January to March 2020. Seeds were densely planted in 15-cell planting trays and thinned after germination to 4-5 plants per cell. Above ground plant material was harvested just before development of inflorescence stems. Plant material was coarsely shredded by hand before adding 100g to 400mL of 10mM MgSO_4_ and autoclaving for 55 minutes. After cooling to room temperature, the medium was filtered through 0.2µm polyethersulfone membrane filters to maintain sterility and remove plant material. The medium was stored in the dark at 4°C. Before being used for culturing, the medium was diluted 1:10 in 10mM MgSO_4_.

### Assembly and culturing of synthetic communities

Fresh bacterial stocks were prepared by first inoculating isolates into 1mL of ALM shaking at 28°C and growing overnight. Next, 100uL of these cultures were used to inoculate 5mL of ALM shaking at 28°C. Once the cultures were visibly turbid, they were divided into 1mL aliquots with sterile DMSO added to a final concentration of 7% as a cryoprotectant. Stocks were stored at −80°C. An aliquot of each stock was used to estimate bacterial cell density through colony counting on ALM plates. To initiate an experiment, stocks were diluted to densities determined by the target initial titer (1×10^6^ cells) of the community and the number of initial members. Diluted stocks were then combined into desired pools (see “experimental design” below) and used to inoculate 600µL of ALM in sterile 1mL deep-well plates, in triplicate. Deep-well plates were covered with sterilized, loosely fitting plastic lids to allow air exchange. Plates were cultured in the dark at 28°C on high-speed orbital shakers capable of establishing a vortex in the deep-well plates to ensure that the cultures were well-mixed. After 24 hours, 6µL of each culture was manually transferred by multi-channel pipette into new plates containing 594µL of fresh ALM. The new plates were immediately returned to the incubator and the day-old plates were stored at −80°C.

### Experimental design

To study invasion across multiple community contexts, we assembled a set of 15 synthetic bacterial communities from a pool of 48 bacterial strains, representing 24 genera (supplementary table 1). The 15 communities were previously characterized (Dooley 2024) and selected to entail a range in final community richness and composition. These communities ranged in initial richness from 8 to 48 strains and were inoculated into the leaf-based medium (ALM). Communities were inoculated at an initially consistent total cell density, with each member at an equal density reduced in proportion to the richness of the pool. We passaged each community into fresh medium (1:100 dilution) every 24 hours up until its invasion treatment (see below) and for 7 days following the invasion (figure 1). We have previously demonstrated that 7 days is sufficient to allow communities in this system to reach an ecologically stable state (21).

**Figure 1-.**
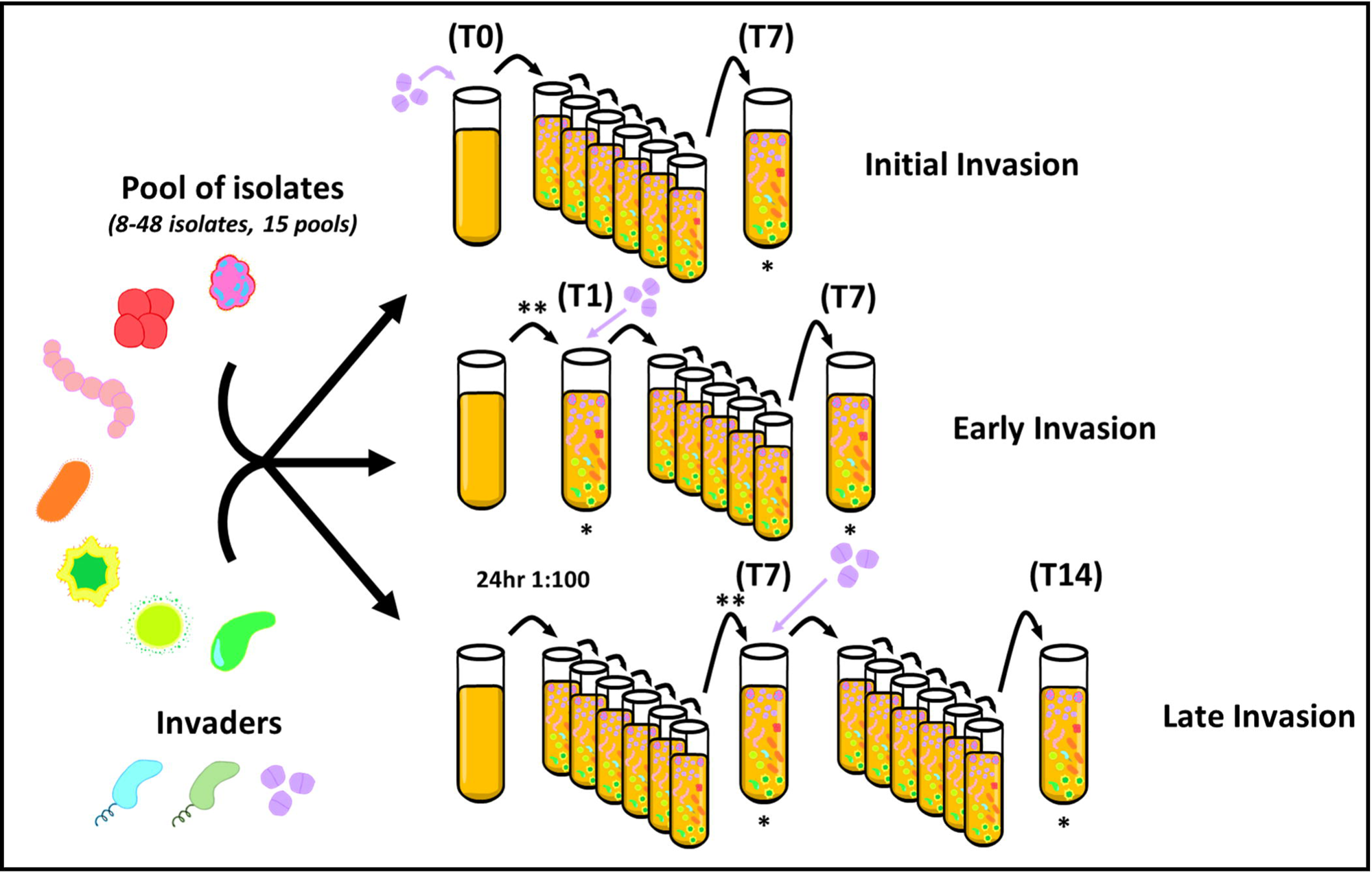
Experimental design: 15 pools of bacterial isolates were separately subjected to invasion by three bacterial isolates across three timing treatments. Across all treatments, passaging always occurred after 24 hours and involved a 1:100 dilution into fresh media. In the “initial invasion” treatment, a given invader was added alongside the other community members at the time of community initiation (T0). In the “early invasion” treatment, a given community was assembled and passaged after 24 hours (T1), before an invader was added immediately after passaging. And in the “late invasion” treatment, a given community was assembled and passaged for 7 days (T7) before an invader was added. Post-invasion, communities were passaged for seven days and characterized using shallow short-read sequencing. Sequenced samples are indicated by (*) while samples filtered for the spent-media assays (methods) are indicated by (**).

To test the effect of invasion timing on community assembly, we invaded each of these 15 communities with three different bacterial invaders at each of three time points in the community assembly process (figure 1). We chose to work with multiple community contexts and invaders to seek general patterns in the effect of invasion timing, rather than outcomes specific to a certain invader and/or community context. We used a strain of *Pseudomonas poae* (Pseudomonas_MEJ082), a strain of *Pseudomonas viridiflava* (Pseudomonas_RMX3.1b), and a strain of *Xanthomonas campestris* (Xanthomonas_S130) as invaders. Each invader was added at a density determined in preliminary experiments to be sufficient to lead to successful invasion in some, but not all, community contexts (“determining appropriate invading densities” below). These densities were 0.1% of estimated total community density for *X. campestris* and 10% for both *P. poae* and *P. viridiflava*.

We assessed three invasion timing treatments that targeted distinct phases of the community dynamics. In the “initial” invasion treatment, invaders were added to the community during initial community assembly (T0), when the assembly process had just begun. In the “early” invasion treatment, invaders were added immediately after the first round of passaging (24 hours, T1), which is an especially dynamic point in the assembly process (21). And in the “late” invasion treatment, invaders were added after 7 days of growth (entailing 6 rounds of passaging, T7), when community dynamics had reached a steady state. After adding invaders, communities were passaged for an additional 7 days. Although the “initial” invasion treatment is not an “invasion” in that the invaders are initially present, it serves as an important reference in defining a baseline of (i) whether a given invader could persist in each community context and (ii) the final community composition. We used NGS to characterize the composition of these communities by mapping short-reads back to previously assembled reference genomes (“sequencing and read mapping” below).

### Determining appropriate invading densities for invaders

Given that we aimed to assess the effect of invaders on the composition of invaded communities, we wanted to avoid instances where invader density was too low to impact a community or so high that an invader would dominate every invaded community. To identify an appropriate density for each invading isolate, 8 of the communities (#8-15) were invaded with each invader across a range of initial densities (0.01%, 0.1%, 1%, 10%, 25%, 100% of estimated invaded community density). These cultures were passaged as described for the main experiment, and invader presence was tracked over time by spot plating 20µL of each culture from each timepoint onto 1X TSA plates containing 70µg/mL gentamicin. The invader isolates had been previously transformed to contain gentamicin-resistance cassettes through mini-Tn7 insertion (22), allowing us to track the presence of the invaders over time. In this way, we identified densities for each invader that produced a variety of invasion outcomes (i.e., failure to establish versus persistence at variable abundances).

### Spent media assays

We performed spent media assays that relate to the “early” and “late” invasion treatments. For the “early” treatment, we isolated spent medium from uninvaded communities after 24 hours of growth (immediately prior to passaging and addition of invaders). For the “late” treatment, we isolated spent medium from uninvaded communities after 7 days of growth (immediately prior to the 7^th^ passage). We isolated spent medium from each community by pelleting the bacterial cells (centrifuged for 10 minutes at 3000 RCF) and filtering ∼150µL of supernatant through 0.2µm polytetrafluoroethylene filtration plates (Pall Corporation, Port Washington, New York, USA). The filtrate was then pooled by community (to produce a representative spent medium for a given community, homogenizing variation among replicates) and amended with M9 salts (at a final concentration of 0.3X) to ensure minimal metabolic needs were met and focus on unused/produced sources of carbon. Prior to inoculation, invader stocks were pelleted and washed in 10mM MgSO_4_ twice to minimize media carryover from the stocks. Each invader was subsequently resuspended in 10mM MgSO_4_, and 5µL was inoculated into 200µL of each spent medium in triplicate and cultured at 28°C in 96-well clear bottom plates. Negative controls were present in each plate, containing only 10mM MgSO_4_ buffer and M9 salts (0.3X). These negative controls were used to subtract background growth from the invaders cultured in spent medium. Growth was assessed by optical density (OD600) after 24 and 48 hours.

### DNA Extraction

DNA was extracted from synthetic communities using an enzymatic digestion and bead-based purification. Cell lysis began by adding 250µL of lysozyme buffer (TE + 100mM NaCl + 1.4U/µL lysozyme) to 300µL of thawed sample and incubating at room temperature for 30 minutes. Next, 200µL of proteinase K buffer (TE + 100mM NaCl + 2% SDS + 1mg/mL proteinase K) was added. This solution was incubated at 55°C for 4 hours and mixed by inversion every 30 minutes. After extraction, the samples were cooled to room temperature before adding 220µL of 5M NaCl to precipitate the SDS. The samples were then centrifuged at 3000 RCF for 5 minutes to pellet the SDS. A Tecan (Männedorf, Switzerland) Freedom Evo liquid handler was used to remove 600µL of supernatant. The liquid handler was then used to isolate and purify the DNA using SPRI beads prepared as previously described (23). Briefly, samples were incubated with 200µL of SPRI beads for 5 minutes before separation on a magnetic plate, followed by two washes of freshly prepared 70% ethanol. Samples were then resuspended in 50µL ultrapure H2O, incubated for 5 minutes, separated on a magnetic plate, and the supernatant was transferred to a clean PCR plate. Purified DNA was quantified using a Picogreen assay (ThermoFisher, Waltham, MA, USA) and diluted to 0.5ng/µL with the aid of a liquid handler.

### Sequencing library preparation

Libraries were prepared using Illumina (San Diego, CA, USA) Nextera XT kits. Our protocol differed from the published protocol in two ways: 1) the tagmentation reaction was scaled down such that 1µL of purified DNA, diluted to 0.5ng/µL, was added to a solution of 1uL buffer + 0.5µL tagmentase, and 2) a KAPA HiFi PCR kit (Roche, Basel, Switzerland) was used to perform the amplification in place of the reagents included in the Nextera XT kit. PCR mastermix (per reaction) was composed of: 3µL 5X buffer, 0.45µL 10mM dNTPs, 1.5µL i5/i7 index adapters, 0.3µL polymerase, and 5.75µL ultrapure H2O. The PCR protocol was performed as follows: 3 minutes at 72 °C; 13 cycles of 95 °C for 10 seconds, 55 °C for 30 seconds, 72 °C for 30 seconds; 5 minutes at 72 °C; hold at 10 °C. Sequencing libraries were manually purified by adding 15µL of SPRI beads and following the previously described approach, eluting into 12µL of ultrapure H2O. Libraries were quantified by Picogreen assay, and a subset of libraries were run on an Agilent 4200 TapeStation system to confirm that the fragment size distributions were of acceptable quality. The libraries were then diluted to a normalized concentration with the aid of a liquid handler and pooled. The pooled libraries were concentrated on a vacuum concentrator prior to size selection for a 300-600bp range on a Blue Pippin (Sage Science, Beverly, MA, USA). The distribution of size-selected fragments was measured by TapeStation. Size-selected pool libraries were quantified by Picogreen assay and qPCR (KAPA Library Quantification Kit).

### Sequencing and read mapping

We characterized the compositions of our synthetic communities with a shallow metagenomics approach. Samples were sequenced on a NovaSeq 6000 platform. Reads were quality filtered and adapter/phiX sequences were removed using BBDuk from the BBTools suite (24). To address ambiguously mapped reads resulting from genomic similarity between some closely related isolates, reads were mapped to reference genomes using Seal (BBTools) twice, once with the “ambig” flag set to “toss” (where ambiguously mapped reads were left out) and once with the “ambig” flag set to “random” (where ambiguously mapped reads were randomly distributed to equally likely references). By comparing the results between these two strategies, we identified sets of reference genomes that resulted in high numbers of ambiguous reads (due to similarity). We corrected for this ambiguity by identifying the number of reads that were removed in the “toss” setting (i.e., the difference in mapped reads between the “toss” and “random” settings) and reallocating those reads based on the proportion of reads unambiguously mapped to each isolate in the “toss” setting (as those proportions represents our best estimate of the true relative abundances of similar isolates). To avoid mischaracterizing the composition of our synthetic communities due to contamination or non-specific mapping, isolates with less than 1% of total mapped reads for a given sample were ignored.

### PERMANOVA analysis

Single-factor PERMANOVA tests were used to determine if the shifts in community composition resulting from the invasion timing treatments were distinct from the uninvaded control communities. Tests were performed using the “adonis2” function from the R package “vegan” (v2.6-4, ref. 25). Bray-Curtis dissimilarity was used to measure the compositional effect of a given treatment. All tests were performed with 999 permutations and the permutations were blocked by community identity.

### Statistical analysis and data visualization

Statistical analysis and figure generation was performed in R v4.0.2 (ref. 26) with aid from the following packages: tidyverse v1.3.0 (ref. 27), reshape2 v1.4.4 (ref. 28), car v3.0-11 (ref. 29), vcd v1.4-11 (ref. 30), and vegan v2.6-4 (ref. 25). All scripts are provided in the supplementary materials.

## Results

### Invasions were commonly unsuccessful, and more so in high richness communities

We define a successful invasion as one in which an invader persists in the community to which they were introduced, regardless of the invader’s relative abundance. In general, successful invasions were uncommon (table 1). Across all invaders and all communities, successful invasion was observed ∼24% of the time. In these instances of successful invasion, the presence of the invader resulted in the exclusion of at least one resident community member 46% of the time. Some invaders were more successful than others (Chi-square test of independence: p-value < 2e^−9^, supplementary figure 1a), with *X. campestris* displaying the highest success rate at 39%, *P. viridiflava* showing the lowest at 6%, and *P. poae* with an intermediate rate of 27% (table 1a). Interestingly, the most successful invader displayed the lowest mean relative abundance (0.03±0.02), while the less successful invaders were more likely to attain a higher relative abundance (mean relative abundance for *P. viridiflava* and *P. poae* were 0.27±0.12 and 0.16±0.18, table 1a).

**Table 1-.** Summary of invasion outcomes by A) invader and B) community. Success rate is calculated as the number of successful invasions out of the total number of invasions.

We also observed a statistical association between invaded community and invasion success, with communities 3, 4 and 6 experiencing notably high invasion success rates across invaders (table 1b, supplementary figure 1b). Given that our communities varied in richness across invasion treatments, we used logistic regression to analyze the relationship between invasion success or failure and richness of an invaded community (supplementary table 2). We found that the probability of successful invasion decreased as the richness of communities prior to invasion increased, but with a low average marginal effect; an increase in richness of one was associated with a 3.3% reduction in the probability of an introduction leading to a successful invasion (figure 2a).

**Figure 2-.**
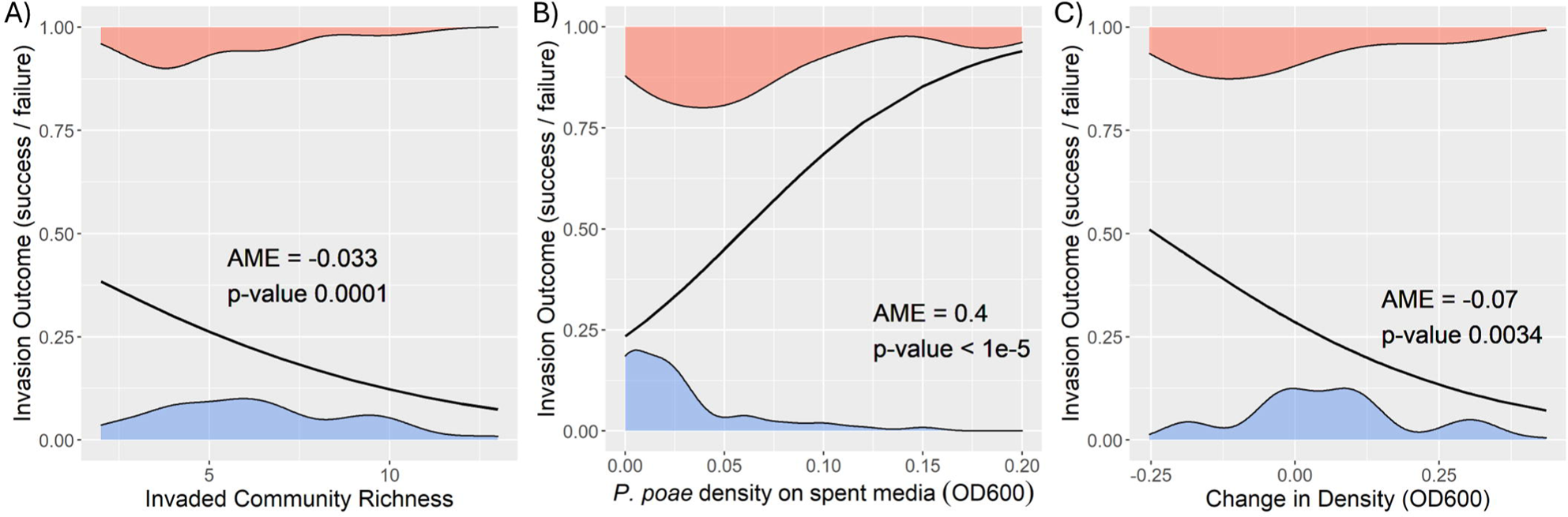
Invaded community richness and estimates of community resource use efficiency were predictive of invasion outcome: Plots depicting the results of logistic regressions analyzing the relationship between invasion outcome (success/failure) with A) the richness of an invaded community B) the growth (optical density) of the invader *Pseudomonas poae* (MEJ082) on spent-media, and C) the change in density between day 1 and day 6 of the “late” invasion communities prior to invasion. Average marginal effects (AME) describe the change in probability of a successful invasion associated with an increase in richness of 1 or a change in density of 0.1. Density plots at the top and bottom of each plot represent the observed distributions of invaded community richness, density, or change in density, partitioned by invasion outcome (blue = failed invasion, red = successful invasion).

### Resource use efficiency was associated with invasion success

If invaders are more successful when introduced into communities with an abundance of metabolites suitable for cross-feeding, then we would expect invasion success to be related to low resource use efficiency. To examine this possibility, we evaluated how well invaders were able to grow on the “spent media” of a community prior to invasion (methods) and tested whether growth on spent media was predictive of invasion outcome. Briefly, we filtered each of the communities after one day and seven days of growth (representing the communities immediately prior to the “early” invasion and “late” invasion treatments, respectively) to remove bacterial cells and isolate sterile “spent” medium. We then assessed the growth of each of the three invader species when cultured on each spent medium, measuring OD600 after 48 hours.

Extensive growth was rare and generally restricted to *P. poae* (supplementary figure 2), which was the invader with an intermediate probability of invasion success at 27%. Although there was insufficient variation in invader growth to explore how community composition influenced the growth of *P. viridiflava* and *X. campestris,* we used logistic regression to analyze the relationship between growth of *P. poae* on spent media from the 15 communities and whether an invasion was successful. We found that a 0.1 increase in *P. poae* optical density was associated with a ∼40% average increase in the probability of a successful invasion (supplementary table 2, figure 2b). Thus, for *P. poae*, invasion success was positively associated with its ability to use the resources unused or produced by a given community.

We additionally wanted to assess if a change in resource use efficiency over the course of community assembly was associated with invasion outcome. To approximate a change in resource use efficiency, we compared the initial (T1) and late stage (T6) densities (OD600) of each uninvaded community. Generally, community density modestly increased over this period, with a statistically significant average increase in optical density of 0.035 (one sample t-test: p-value 0.01). We used logistic regression to relate these changes in density over time with invasion outcome in the late-invasion treatment and observed a significant negative relationship; namely, a decrease in density of 0.1 during assembly was associated with a 7% increase in the probability of a successful invasion (supplementary table 2, figure 2c). This did not simply reflect that less dense communities were more prone to invasion but rather that communities that declined most extensively were more vulnerable to invasion; this was clear because pre-invasion density itself was not significantly associated with invasion success by logistic regression (p-value 0.52).

### Invaders had an outsized effect on the most abundant members of invaded communities

We observed wide variation in how community compositions shifted after invasion, successful or otherwise. In some instances of successful invasion, the change in community composition could largely be attributed to the presence of the invader. However, there were also instances where, despite occupying a low relative abundance or even failing to persistently invade (figure 3a), an invader had a large effect on community composition. There is debate as to whether common and rare species suffer different vulnerabilities to invasion (31–34). Thus, we asked whether rare or common species were more greatly impacted by the invasion of new species by calculating the log-ratio of post-invasion and pre-invasion relative abundances for each community member that persisted in a given invaded community. We then asked how those log-ratios related to the rank abundance of each community member, with a total of 9 ranks set by the maximum observed final richness. We observed that the greatest log-ratio change in relative abundance was associated with the community members with the highest rank-abundance in the pre-invasion community (one-way ANOVA between ranks: F_8,507_ = 20.69, p-value < 2e^−16^, followed by Tukey’s honest significance test) (figure 3b). Indeed, only the highest rank abundance community members showed a significant decrease in relative abundance (one-sample t-test: p-value < 2.2e^−16^), while ranks 2-7 showed a significant increase (p-values < 0.05) and ranks 8/9 showed no significant change. Thus, across our full set of invasion outcomes, we observed a surprisingly consistent pattern in which the most abundant species in the community were most affected by the invader.

**Figure 3-.**
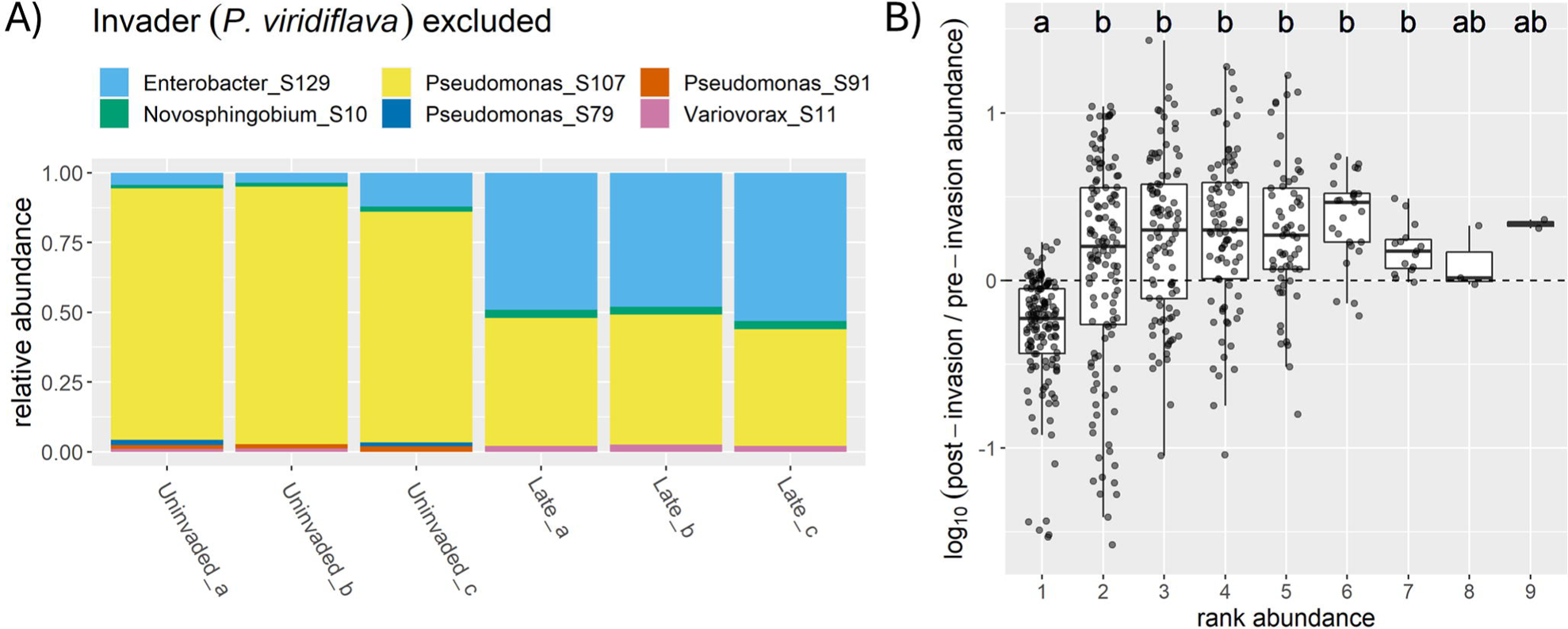
Highest rank abundance community members were most affected by invasion: A) Example demonstrating an instance where the invader (*P. viridiflava*) had a strong effect on the most-abundant member of the invaded community (Pseudomonas_S107), despite not persisting in the community. B) Bar and whisker plots display the distribution of log-ratios between post- and pre-invasion relative abundances for all community members across all communities, grouped by rank abundance in the respective uninvaded context (rank 1 = most abundant). Significant differences between groups determined through Tukey’s honest significance test.

### Effect of invasion timing can be explained by invader growth on spent media

To assess the effect of invasion timing on invasion outcome, we first compared the probability of a successful invasion across invasion treatments. A Chi-square test of independence revealed a significant association between invasion treatment and invasion outcome (success/failure) (p-value < 5e^−5^), with successful invasions more common than expected in the early invasion treatment and less common than expected in the initial invasion treatment (figure 4), when all invaders were considered jointly. The occurrence of successful invasions in the late invasion treatment did not significantly deviate from the null expectation. To test whether invasion timing had an impact on the composition of communities, we calculated the average Bray-Curtis dissimilarity between samples of each timing treatment and uninvaded control communities, disregarding the invader if it was present. We then used pairwise single-factor PERMANOVA tests to determine if the invasion treatments resulted in community compositions distinct from the uninvaded controls (methods). We observed that all invasion timing treatments resulted in significantly distinct community compositions (all p-values ≤ 0.003 after multiple testing correction).

**Figure 4-.**
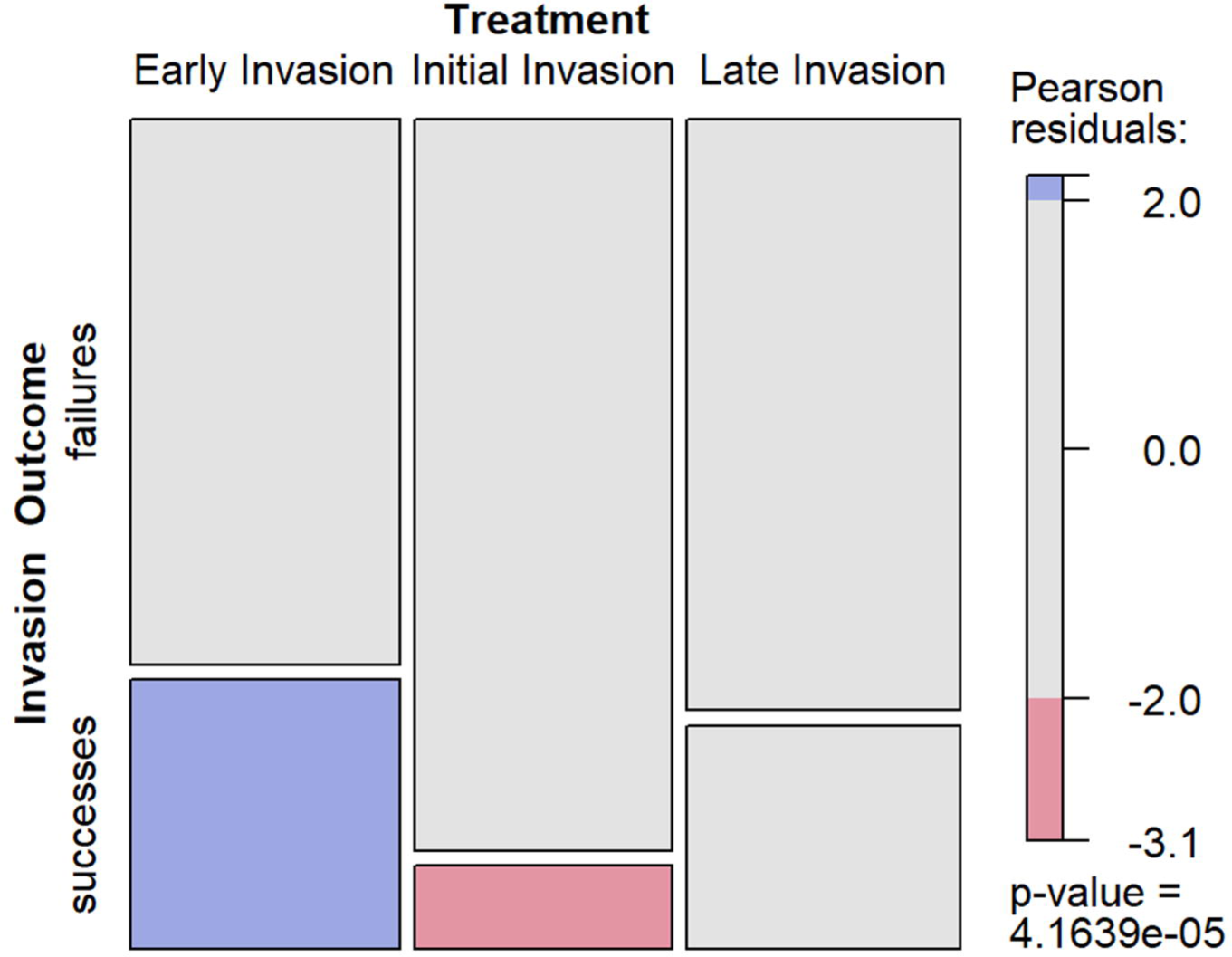
Invasion timing affected invasion outcome: Mosaic plot representing the relative frequencies of invasion outcome across the three invasion timing treatments. Invasion outcome was defined as a “success” (bottom box) if the invader persisted over time, or a “failure” (top box) if the invader was ultimately excluded. Chi-square test of independence demonstrates a significant association between invasion outcome and invasion timing (p-value 4.16e^−5^), with the Pearson residuals indicating invasion success was significantly over-represented in the early-invasion treatment (highlighted in blue) but under-represented in the initial-invasion treatment (highlighted in red). The late-invasion results did not significantly deviate from the null expectations. Pearson residuals >2 or <-2 indicate a count value greater than 2SD from the expectation.

We next sought to determine if there were differences in the extent to which invasion timing treatments affected community composition by comparing the Bray-Curtis dissimilarities calculated for each treatment relative to the uninvaded control. A one-way ANOVA found that the average Bray-Curtis dissimilarity differed between the invasion treatments (F_2,381_ = 5.23, p-value 0.006, supplementary table 3). Post-hoc tests showed strong support for a significant difference in dissimilarity between the early and initial invasion treatments with an average difference of 0.08 (Tukey’s honest significant test: 95% CI [0.02, 0.14], p-value = 0.005, supplementary table 3). There was also marginal statistical support for a difference between the early and late invasion treatments (Tukey’s honest significant test: 95% CI [-0.005, 0.11], p-value = 0.081, supplementary table 3).

Given that we had previously identified associations between invasion outcome and both community richness and invader growth on spent media, and that these factors may change over the course of community assembly, we included these variables as covariates in additional analyses (ANOVA) to determine if they could explain the effect of invasion timing (supplementary table 4). Controlling for differences in richness did not strongly affect the significance of invasion timing in these models, demonstrating that this factor could not explain the effect of timing. However, invader growth on spent media was a significant covariate and, importantly, reduced the effect of invasion timing. Note, however, that this analysis only considered the “early” and “late” invasion treatments, as those were the only treatments for which we could measure growth on spent media prior to invasion.

## Discussion

We hypothesized that the timing of an invasion would impact its outcome because factors affecting ecological invasions change over the course of community assembly. Community diversity is one such dynamic factor. Indeed, the relationship between diversity and community invasibility has long been studied in plant (11,13,15,35,36) and experimental bacterial communities (10,12,14), as well as in the context of enteric pathogens (37,38). Work in multiple biological systems has shown that invasion is less successful in diverse communities (10–12,14). This phenomenon may be explicable since increased diversity can result in more complete occupancy of available niche space, thus increasing community resource use efficiency and reducing resources that are available for an invader (10,15,39). Our results are in general alignment with this effect, less diverse communities were more likely to be successfully invaded (figure 2a) and, at least for *P. poae*, growth on spent medium was positively associated with invasion success (figure 2b). In addition, the observed effect of invasion timing was reduced when we incorporated invader growth on spent media (supplementary table 4), suggesting that the impact of invasion on community composition was, at least in part, modulated by available resources.

Another relevant factor is community composition, which inherently changes as community members are filtered out during assembly. This is relevant to invasibility in that invaders can be excluded if they compete with species that share similar nutrient requirements (40–42). This mechanism, referred to as a “sampling effect”, is also related to richness, as higher richness increases the chance that a community contains species capable of excluding an invader through competition (15–17). Thus, as communities settle into stable composition of reduced complexity during the assembly process, invasibility, as mediated through this mechanism, should increase.

We can use the effects described above to interpret the differences we observed between invasion timing treatments; namely, why the early invasions were most susceptible to invasion (figure 4). First, the initial invasion treatment was less prone to invasion most likely because the increased richness encountered by invaders in this treatment (as no competitive exclusion could yet have occurred) increased the chances that a community member could exclude the invader, thus decreasing the chance of successful invasion through a “sampling effect”. On the other hand, the enhanced resistance to invasion in the late treatment was associated with decreased invader growth on spent media (supplementary table 4). One contributing factor to the decrease in resources available in the spent media was likely the increase in community density (figure 2c). Why then were communities in the early treatment prone to invasion? We hypothesize that resource use efficiency in the early treatment, while community dynamics were still transient, may have been depressed because of strong early competition (19,20) that eased over time as assembly continued and competitors were excluded. Additionally, communities may have achieved higher resource use efficiency over time through mechanisms such as changes in metabolic regulation or evolution that resulted in community members becoming better able to utilize available resources (43–46).

Consistent with our results, Rivett et al. (2018) reported decreased success of invasions later in the assembly process and identified change in resource availability across community assembly as a mechanism underlying invasion outcome. In that study, assembly occurred in a static environment with no nutrient replenishment. Our method of assembly through passaging represents a distinct assembly process akin to environments with higher sustained metabolic activity resulting from periodic influxes of resources (e.g., the gut environment). Despite the difference in nutrient dynamics between our two systems, the convergence of our results suggests the relationship between invasion timing and outcome is robust across environments.

Previous work investigating the importance of invasion timing has focused on the synchronization of invasions with periods of increased resource availability (16,48). This is also related to work investigating the relationship between disturbance and invasibility, which has posited that disruptions in community resource use efficiency enhances the opportunity for invasion (49–51). Our results are in general alignment with these perspectives, as they rely on periods of increased relative resource availability as predictors of invasion outcome, but in contrast to past work relies on natural, dynamical properties of community assembly rather than extrinsic perturbations. Namely, our work demonstrates that transient dynamics, whether caused through ecological perturbation or the natural assembly process, facilitate invasion.

Interestingly, we observed that even unsuccessful invasions could have large effects on the final composition of the communities from which they were excluded (figure 3a). This finding supports results from Amor et al. (2020), which reported that transient invasion could cause a transition between alternative stable states in a two-member bacterial community and a moderately more complex community derived from soil (52). Our work furthers this result by demonstrating that such a phenomenon might be common, should an invasion occur when a community is still in the transient phase of assembly. More generally, it has been shown that priority effects (53,54) and minor differences in transient states early in the assembly of bacterial communities can lead to divergent final community compositions (43). This context can help us understand why the initial and early invasion treatments led to such distinct community compositions (supplementary table 3) as the communities at T0 and T1 were sufficiently distinct that it is unsurprising that disruption via invasion drove distinct paths of further assembly. This perspective further highlights the importance of transient community states during the assembly process.

Additionally, we observed that it was generally the most abundant member of a community that was most affected by invasion (figure 3b). This finding is relevant from an applied perspective, suggesting that higher abundance members of a community might be more likely to suffer a greater impact from invasion and thus harder to maintain at a consistent relative abundance within an engineered community.

Overall, we show that invasion of a synthetic bacterial community at different points of the community assembly process affects invasion success and the impact of the invaders on the resident community. We found rather consistent increases in the success rate and impact of invasions that occurred during the transient phase of the community assembly process, which aligned with the expected effects of decreased diversity and resource use efficiency. Additionally, common rather than rare community members were most impacted by invaders. And, interestingly, we found strong effects of introduced species that failed to persist on community composition. These results further our understanding of the factors affecting the invasibility of microbial communities, with implications relevant to human health, agriculture, and industry.

## Supporting information

Rcode and associated data files

## Acknowledgements

This work was supported by the Hutchinson fund at The University of Chicago, the Simons Foundation, and ERC Synergy grant, PATHOCOM (951444, J.B.).

## Data Availability Statement

The datasets generated during and/or analyzed during the current study are available in the NCBI Whole Genome and Sequence Read Archive repositories (accessions: PRJNA953780, PRJNA1073231).

