## Supplementary Figures and Tables for "The repercussions of timing in the invasion of synthetic bacterial communities"

*Supplementary Table 1: isolate details (attached .csv)*

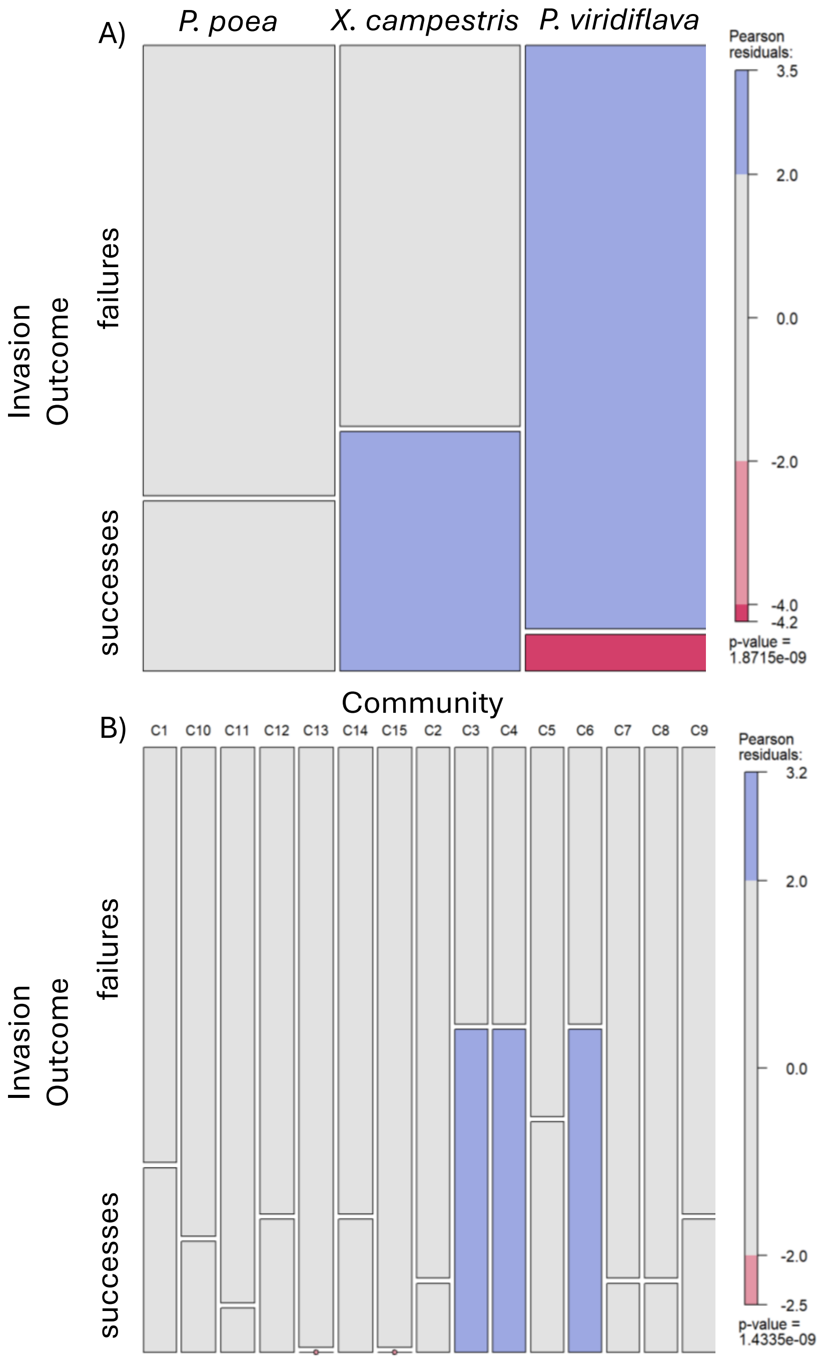


Supplementary figure 1 – Invasion success varied between invader and community: A) Mosaic plot representing successful and failed invasions for each invader. Invasion outcome was defined as a “success” (bottom box) if the invader persisted over time, or a “failure” (top box) if the invader was ultimately excluded. Chi square test of independence shows invader identity was statistically associated with invasion success (p-value 1.87e$^{-9}$). Successful invasions were observed more often than expected for *X. campestris* (highlighted in blue). *P. viridiflava* demonstrated more failures than expected and fewer successes (highlighted in red and blue, respectively. B) Mosaic plot representing successful and failed invasions in each community. Invasion outcome was defined as a “success” (bottom box) if the invader persisted over time, or a “failure” (top box) if the invader was ultimately excluded. Chi square test of independence shows community identity was statistically associated with invasion success (p-value 1.43e$^{-9}$). Successful invasions were more common than expected for communities 3, 4, and 6 (highlighted in blue), and less common than expected for communities 13 and 15 (represented as red dots). Pearson residuals >2 or <-2 indicate a count value greater than 2SD from the expectation.

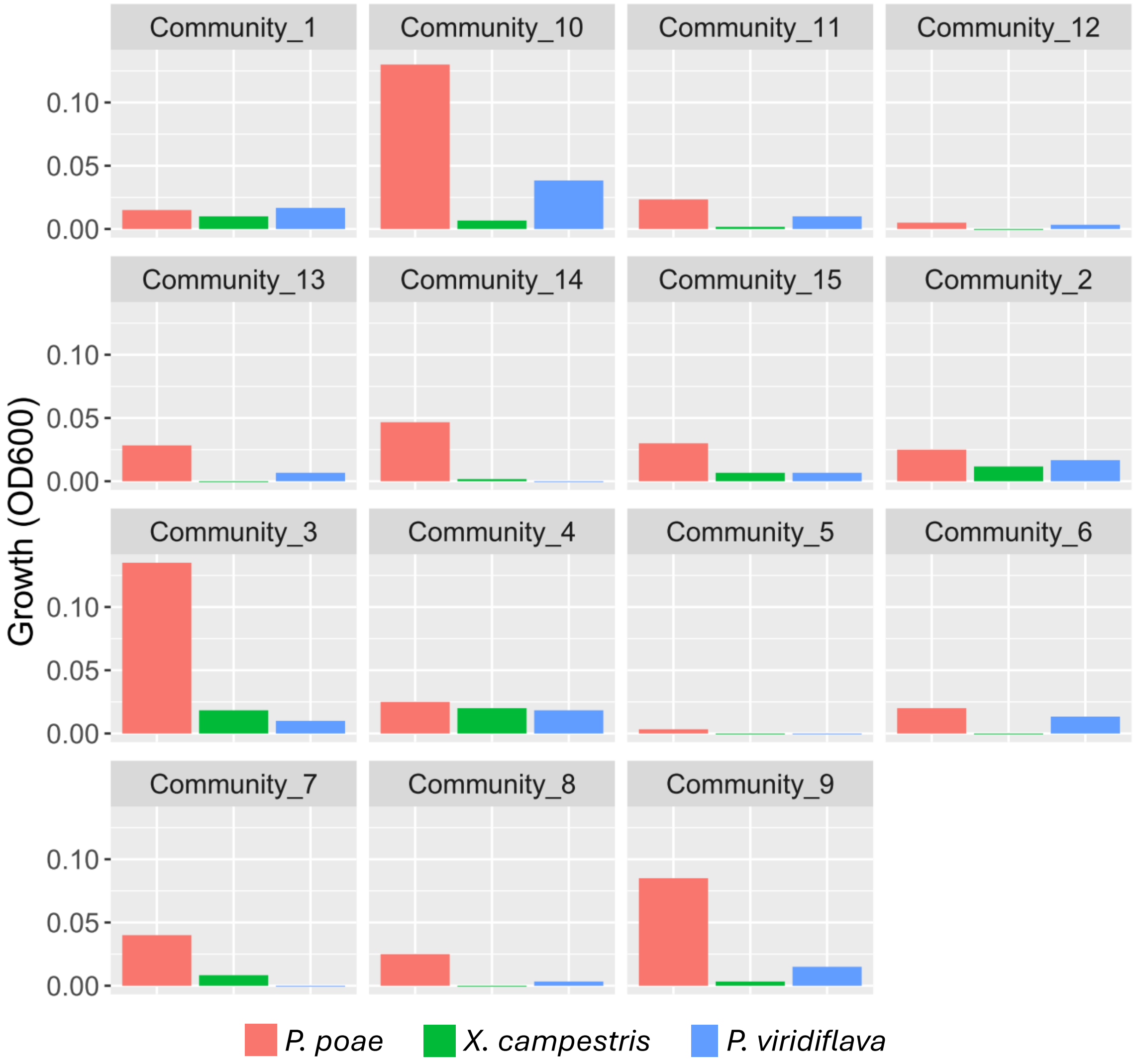


Supplementary figure 2 – Growth on spent media varied by community and invader: Bar plots display the average optical density, as a proxy for growth, of each invader across spent-medium from each community. Averages were taken across both the initial and early invasion treatment spent media.

| Logistic Regression | | | | |
|---|---|---|---|---|
| **A) invasion outcome ~ richness** | | | | |
| **Variable** | **Estimate** | **Std. Error** | **z value** | **Pr > \|z\|** |
| (intercept) | -0.098 | 0.301 | -0.325 | 0.745 |
| Richness | -0.187 | 0.051 | -3.66 | 0.0003 |
| Null Deviance | 433.5 | on 394 df | | |
| Residual Deviance | 418.74 | on 393 df | | |
| **B) invasion outcome ~ density on spent media (*P. poae*)** | | | | |
| (intercept) | -1.18 | 0.33 | -3.58 | 0.0003 |
| Density | 1.96 | 0.6 | 3.25 | 0.001 |
| Null Deviance | 121.91 | on 89 df | | |
| Residual Deviance | 107.46 | on 88 df | | |
| **C) invasion outcome ~ change in density (day 1 vs day 6)** | | | | |
| (intercept) | -0.918 | 0.199 | -4.61 | $< 4e^{-6}$ |
| Change in Density | -0.379 | 0.141 | -2.69 | 0.007 |
| Null Deviance | 158.56 | on 134 df | | |
| Residual Deviance | 150.48 | on 133 df | | |

| D) Average Marginal Effects | | | | | ***95% CI*** | |
|---|---|---|---|---|---|---|
| **Variable** | **AME** | **SE** | **z** | **p-value** | **lower** | **upper** |
| Richness | -0.033 | 0.009 | -3.82 | 0.0001 | -0.05 | -0.016 |
| Density (*P. poae*) | 0.402 | 0.095 | 4.22 | 0.0000 | 0.216 | 0.589 |
| Change in Density | -0.071 | 0.024 | -2.93 | 0.0034 | -0.118 | -0.024 |

Supplementary Table 2 – Invaded community richness and resource use efficiency were associated with invasion success: Details of logistic regressions analyzing the relationships between invasion outcome with A) the richness of an invaded community (“Richness”), B) the density of a specific invader (P. poae) on spent-media (“Density”), and C) the change in community density between day 1 and day 6 (all density measurements represent optical density at 600nm). D) The average marginal effects of: an increase of 1 in richness, an increase of 0.1 in the density of P. poae, and a change in density of 0.1 are presented in the lower section of the table.

| A) One-way ANOVA: Bray-Curtis dissimilarity ~ invasion treatment | | | | |
|---|---|---|---|---|
| ***variable*** | ***df*** | ***sum of squares*** | ***F*** | ***p-value*** |
| invasion treatment | 2 | 0.456 | 5.23 | 0.006 |
| residuals | 381 | 16.6 | | |

| B) Tukey's Honest Significance Test: invasion treatment | | | | |
|---|---|---|---|---|
| **invasion treatment** | **mean difference** | **95% CI lower** | **95% CI upper** | **adjusted p-value** |
| initial – late | -0.026 | -0.088 | 0.036 | 0.596 |
| early – late | 0.056 | -0.005 | 0.117 | 0.081 |
| early – initial | 0.082 | 0.021 | 0.143 | 0.005 |

Supplementary Table 3 – The initial invasion treatment was the most dissimilar treatment relative to the uninvaded communities: A) One-way ANOVA (Type III) results analyzing the relationship between Bray-Curtis dissimilarity (relative to the uninvaded communities) and invasion timing treatment. B) Tukey's Honest Significance Test results comparing the differences in Bray-Curtis dissimilarities between the invasion timing treatments.

| A) One-way ANOVA: Bray-Curtis dissimilarity ~ invasion treatment | | | | |
|---|---|---|---|---|
| ***variable*** | ***df*** | ***sum of squares*** | ***F*** | ***p-value*** |
| invasion treatment | 2 | 0.456 | 5.23 | 0.006 |
| residuals | 381 | 16.6 | | |

| B) One-way ANCOVA: Bray-Curtis dissimilarity ~ invasion treatment + richness | | | | |
|---|---|---|---|---|
| invasion treatment | 2 | 0.456 | 5.22 | 0.006 |
| richness | 1 | 0.015 | 0.337 | 0.562 |
| residuals | 380 | 16.6 | | |

| C) One-way ANCOVA: Bray-Curtis dissimilarity ~ invasion treatment + spent media growth | | | | |
|---|---|---|---|---|
| invasion treatment | 1 | 0.202 | 5.141 | 0.024 |
| spent media growth | 1 | 0.324 | 8.24 | 0.004 |
| residuals | 254 | 9.99 | | |

Supplementary Table 4 – Invader growth on spent media is a significant covariate in the relationship between invasion timing and outcome: A) One-way ANOVA results from supplementary table 3, for reference. B) One-way ANCOVA including pre-invasion community richness as a covariate. C) One-way ANCOVA including invader growth on spent media as a covariate (limited to “early” and “late” invasion treatments).
