## Supplementary material for "The repercussions of timing in the invasion of synthetic bacterial communities": Rcode and associated data files: supplementary_materials.R

```
library(tidyverse)
library(reshape2)
library(vcd)
library(margins)
library(generics)
library(car)
library(vegan)

setwd("...")

### treatment key ###
# Tr1 = no invader, T7
# Tr2 = invader added at T7 (late invasion)
# Tr3 = invader added at T0 (initial invasion)
# Tr4 = no invader, T1
# Tr5 = invader added after T1 (early invasion)


# load the data
################################################################################
data = read.csv("invasion_readcounts_corrected.csv", header = TRUE, row.names = 1)
metadata = read.csv("invasion_metadata_corrected_plus.csv", header = TRUE, row.names = 1)
metadata$sample = gsub("_T1_rc","", metadata$sample) %>% substr(.,1,nchar(.) -1)
metadata$name = paste(metadata$sample, metadata$rep, sep = "")
colnames(data) = metadata$name
taxonomy = read.csv("taxonomy_invasion.csv", header = TRUE)
rownames(taxonomy) = taxonomy$isolate
################################################################################

# prune some of the data
################################################################################
# make relative abundance data
relabund = apply(data, 2, function(x){x / sum(x)})

# prune by low relative abundance (< 1%)
relabund[which(relabund < 0.01)] = 0
relabund = apply(relabund, 2, function(x){x / sum(x)}) # re-normalize

# remove the negative controls
relabund = relabund[,-which(metadata$community == "NC")]
metadata = metadata[-which(metadata$community == "NC"),]

# prune some samples that seem to have a systematic error
# Treatment 3 communities 1-8 (plate 5 of the experiment)
relabund = relabund[,-which(metadata$treatment == "Tr3" & metadata$invader == "I4" & metadata$rep == "b" &
                              as.numeric(substr(metadata$community, 2, nchar(metadata$community))) < 9)]
metadata = metadata[-which(metadata$treatment == "Tr3" & metadata$invader == "I4" & metadata$rep == "b" &
                             as.numeric(substr(metadata$community, 2, nchar(metadata$community))) < 9),]

# prune the data so that only isolates that were detected in at least 2 replicates are present in the data
test = relabund
for (i in 1:length(unique(metadata$sample))){
  
  subset = relabund[,grep(unique(metadata$sample)[i], colnames(relabund))]
  
  if (length(subset) > (length(relabund[,1]))) { # this IF & for-loop deals with the fact that samples without replicates throw an apply error
    detected = apply(subset, 1, function(x){sum(x>0)}) >= 2
    
    for (i in grep(unique(metadata$sample)[i], colnames(relabund))){
      test[,i] = as.numeric(relabund[,i]) * detected
    }  
  }
}
relabund = test
rm(subset,detected,i, test)
relabund = apply(relabund, 2, function(x){x / sum(x)}) # re-normalize
################################################################################

# remove S. panni
################################################################################
# the S_panni invader was never detected, which suggests to me that my reference genome was junk and/or the invader was not actually
# a sphingomonas isolate. So, we removed it from the analysis.
relabund = relabund[,-which(metadata$invader == "I3" & metadata$treatment != "Tr1" & metadata$treatment != "Tr4")]
metadata = metadata[-which(metadata$invader == "I3" & metadata$treatment != "Tr1" & metadata$treatment != "Tr4"),]
################################################################################

# make some richness metadata
################################################################################
richness = read.csv("richness_meta.csv", header = TRUE)

temp = relabund[,grep("Tr1", colnames(relabund))]
temp = data.frame(name = colnames(temp), n = apply(temp,2,function(x){length(which(x > 0))})) %>%
  merge(., metadata, by = "name")
temp = temp %>% group_by(community) %>% summarize(mean_n = mean(n))
richness = merge(richness,temp, by = "community")
rm(temp)
metadata$avg_n = sapply(metadata$community, function(x){richness$mean_n[richness$community == x]})

# Don't always want to use average richness, because for each Early and Late invasion (and the initial), we know the richness of the invaded community
kev_func = function(df){apply(df, 2, function(x){length(which(x > 0))})}
rich = metadata$name[-grep("Tr1|Tr4", metadata$name)] %>% lapply(., function(x){
  #print(x)
  temp = if(substr(x,1,3) == "Tr2"){ # if Tr2 (late), find the average richness of the appropriate Tr1 samples 
    relabund[,grep(gsub("Tr2_","",x), metadata$name[which(metadata$treatment == "Tr1")], value = TRUE)] %>% 
      as.data.frame %>% kev_func() %>% mean() } else {
        if(substr(x,1,3) == "Tr3"){ # if Tr3 (initial), find the T0 richness
          richness$Pool_n[metadata$community[which(metadata$name == x)] == richness$community]
        } else { # if Tr5 (early), find the Tr4 (T1) richness
          relabund[,grep(gsub("Tr5_","",x), metadata$name[which(metadata$treatment == "Tr4")], value = TRUE)] %>% 
            as.data.frame %>% kev_func() %>% mean()
        } }
  data.frame(name = x, richness = temp)
}) %>% do.call(rbind.data.frame,.)
rich = rich[!is.na(rich$richness),]

################################################################################

# the invaders
################################################################################
invaders = relabund[c("MEJ082","LMC-P-011_S130","RMX3_1b"),] %>% t()

# reformat the data
invaders = melt(invaders)
colnames(invaders) = c("name","invader","relabund")  
invaders$num = sapply(invaders$invader, function(x){taxonomy$invader[grep(x, taxonomy$isolate)]})
invaders = invaders[sapply(1:nrow(invaders), function(x){length(grep(invaders$num[x], invaders$name[x])) > 0}),]

# add metadata
invaders = merge(invaders, metadata, by = "name")
invaders = invaders[-which(invaders$treatment == "Tr1" | invaders$treatment == "Tr4"),]
invaders = invaders[,c("name","sample","invader.x","invader.y","treatment","community","relabund")]
invaders = merge(invaders,richness, by = "community")
invaders$success = sapply(invaders$relabund, function(x){ifelse(x > 0, 1, 0)})
invaders$treatment[which(invaders$treatment == "Tr3")] = "Initial Invasion"
invaders$treatment[which(invaders$treatment == "Tr2")] = "Late Invasion"
invaders$treatment[which(invaders$treatment == "Tr5")] = "Early Invasion"
invaders = merge(invaders, rich, by = "name")

# how successful were invaders altogether?
sum(invaders$success)/nrow(invaders)

# who was the most common?
table(invaders$invader.x[which(invaders$relabund > 0)])

# who reached the highest average abundance?
invaders[which(invaders$relabund > 0),] %>% group_by(invader.x) %>% summarise(mean = mean(relabund), sd = sd(relabund))

# how often did each invader invade? (Supplementary Figure 1a)
table(invaders$invader.x, invaders$success)[,2]/table(invaders$invader.x) # number successful invasions / total attempts
summary(table(invaders$invader.x, invaders$success))
mosaic(~ invader.x + success, data = invaders, direction = c("v","h"), shade = TRUE)

# were some community contexts more likely to be invaded? (Supplementary FIgure 1b)
table(invaders$community, invaders$success)[,2]/table(invaders$community) # number successful invasions / total attempts
summary(table(invaders$community, invaders$success))
mosaic(~ community + success, data = invaders, direction = c("v","h"), shade = TRUE)


### did spent-media growth predict invasion success?
# read in the spent-media data
spent = read.csv(file = "invasion_spent_media.csv", header = TRUE)
spent$growth[which(spent$growth < 0)] = 0
spent = spent[-which(spent$community == "Control"),]
spent = merge(invaders, spent[,c("name","growth")], by = "name")


# Supplementary Figure 2
kev_plot = spent %>% group_by(.,community, invader.x) %>% summarise(mean = mean(growth), sd = sd(growth))
kev_plot$community = gsub("C","Community_", kev_plot$community)
#pdf(file = "./Supplementary_figure2.pdf")
ggplot(data = kev_plot, aes(x = invader.x, y = mean, fill = invader.x)) + geom_col() + facet_wrap(~community, scales = "free_x") +
  theme(axis.text.x = element_blank(), axis.ticks.x = element_blank(), legend.title = element_blank(), 
        text = element_text(size = 15), legend.position = "bottom") + ylab("Growth (OD600)") + xlab("")
#dev.off()


# Figure 2a and Supplementary Table 2 - was invaded community richness associated with invasion outcome?
kev_model = glm(data = invaders, formula = success ~ richness, family = "binomial")
summary(kev_model)
margins(kev_model) %>% summary()
kev_plot = augment(kev_model, type.predict = "response")
kev_plot = kev_plot %>% mutate(invade = ifelse(success == 1, richness, NA), no_inv = ifelse(success == 0, richness, NA))

#tiff("Figure2a.tiff", units="in", width=5, height=5, res=300)
ggplot(kev_plot, aes(x = richness)) + geom_line(aes(y = .fitted), linetype = 1, size = 1) + xlab("Invaded Community Richness") +
  ylab("Invasion Outcome (success / failure)") +
  geom_density(aes(x = no_inv, y = ..scaled../10), na.rm = TRUE, fill = "cornflowerblue", alpha = 0.5) +
  geom_density(aes(x = invade, y = -1*..scaled../10), na.rm = TRUE, position = position_nudge(y = 1), 
               fill = "tomato", alpha = 0.5) + 
  geom_text(x = 5.5, y = 0.4, label = "AME = -0.033\np-value 0.0001", hjust = 0, size = 6) +
  theme(text = element_text(size = 16))
#dev.off()

# Figure 2b and Supplementary Table 2 - is growth on spent media a significant predictor of a successful invasion?
spent$temp = spent$growth*10 # adjust OD600 measurements so that we're on a more reasonable scale for reference
kev_model = glm(data = spent, formula = "success ~ temp", family = "binomial")
kev_model = glm(data = spent[which(spent$invader.x == "MEJ082"),], formula = "success ~ temp", family = "binomial")
kev_model = glm(data = spent[which(spent$invader.x == "RMX3_1b"),], formula = "success ~ temp", family = "binomial")
kev_model = glm(data = spent[which(spent$invader.x == "LMC-P-011_S130"),], formula = "success ~ temp", family = "binomial")
summary(kev_model)
margins(kev_model) %>% summary()

  # diagnostics
  cor.test(spent$temp[which(spent$invader.x == "MEJ082")], predict(kev_model, type = "response"))
  library(DHARMa)
  test = simulateResiduals(fittedModel = kev_model)
  plot(test)

kev_model = glm(data = spent[which(spent$invader.x == "MEJ082"),], formula = "success ~ temp", family = "binomial")
kev_plot = augment(kev_model, type.predict = "response")
kev_plot = kev_plot %>% mutate(invade = ifelse(success == 1, temp, NA), no_inv = ifelse(success == 0, temp, NA))

#tiff("Figure2b.tiff", units="in", width=5, height=5, res=300)
ggplot(kev_plot, aes(x = temp/10)) + geom_line(aes(y = .fitted), linetype = 1, size = 1) + 
  xlab(expression(italic("P. poae")~density~on~spent~media~(OD600))) +
  ylab("Invasion Outcome (success / failure)") +
  geom_density(aes(x = no_inv/10, y = ..scaled../5), na.rm = TRUE, fill = "cornflowerblue", alpha = 0.5) +
  geom_density(aes(x = invade/10, y = -1*..scaled../5), na.rm = TRUE, position = position_nudge(y = 1), fill = "tomato", alpha = 0.5) + 
  geom_text(x = 0.125, y = 0.25, label = "AME = 0.4\np-value < 1e-5", hjust = 0, size = 6) +
  theme(text = element_text(size = 16))
#dev.off()

# Figure 2c and Supplementary Table 2 - is invasion outcome associated with change in community density between beginning (T0) and end (T6) of assembly?
spent = read.csv(file = "invasion_spent_media3.csv", header = TRUE)
spent = spent[-which(spent$community == "Control"),]
spent = merge(invaders, spent[,c("name","preinv","delta6","delta7")], by = "name")

# on average, did communities increase in density?
t.test(spent$delta6)

# logistic regression
kev_model = glm(data = spent, formula = "success ~ preinv", family = "binomial") # preinvasion density isn't significant

spent$temp = spent$delta6*10 # adjust OD600 measurements so that we're on a more reasonable scale for reference
kev_model = glm(data = spent, formula = "success ~ temp", family = "binomial")
summary(kev_model)
margins(kev_model) %>% summary()

kev_model = glm(data = spent, formula = "success ~ delta6", family = "binomial")
kev_plot = augment(kev_model, type.predict = "response")
kev_plot = kev_plot %>% mutate(invade = ifelse(success == 1, delta6, NA), no_inv = ifelse(success == 0, delta6, NA))

#tiff("Figure2c.tiff", units="in", width=5, height=5, res=300)
ggplot(kev_plot, aes(x = delta6)) + geom_line(aes(y = .fitted), linetype = 1, size = 1) + xlab("Change in Density (OD600)") +
  ylab("Invasion Outcome (success / failure)") +
  geom_density(aes(x = no_inv, y = ..scaled../8), na.rm = TRUE, fill = "cornflowerblue", alpha = 0.5) +
  geom_density(aes(x = invade, y = -1*..scaled../8), na.rm = TRUE, position = position_nudge(y = 1), fill = "tomato", 
               alpha = 0.5) + theme(text = element_text(size = 16)) +
  geom_text(x = 0.15, y = 0.3, label = "AME = -0.07\np-value 0.0034", hjust = 0, size = 6)
#dev.off()


# Figure 4 - were invaders more successful in some timing treatments than others?
table(invaders$treatment[which(invaders$success == 1)]) / table(invaders$treatment)
table(invaders$treatment, invaders$success)

#pdf("Figure4.pdf", width = 6, height = 6)
temp = invaders[,c("treatment","success")]
colnames(temp) = c("Treatment","Invasion_Outcome")
mosaic(~ Treatment + Invasion_Outcome, data = temp, direction = c("v","h"), shade = TRUE, 
       set_labels = list(Invasion_Outcome = c("failures","successes")))
#dev.off()  

rm(kev_model,temp)
################################################################################

# rank abundance shifts
################################################################################
# gonna work with an averaged summary
avg_invaders = lapply(unique(invaders$sample), function(x){
  data.frame(
    invaders[grep(x, invaders$sample),][1,c("community","name","sample","invader.x","invader.y","treatment","Pool_n","mean_n")],
    relabund = mean(invaders[grep(x, invaders$sample),"relabund"]),
    success = mean(invaders[grep(x, invaders$sample),"success"]) )
}) %>% do.call(rbind.data.frame,.) 

# gonna need an average of the relative abundance data
avg_rel = lapply(unique(metadata$sample), function(x){
  if(ncol(as.data.frame(relabund[,grep(x, colnames(relabund))])) > 1){ # if there are replicates...
    rowMeans(relabund[,grep(x, colnames(relabund))])} else { # take the average of each row
      relabund[,grep(x, colnames(relabund))] } # and if not, just use the single sample
}) %>% do.call(cbind.data.frame,.)
colnames(avg_rel) = unique(metadata$sample)
avg_rel = as.matrix(avg_rel)

### rank abundance shift - between the compared contexts, for each isolate in the uninvaded T7 context, how does it change?
results = lapply(avg_invaders$sample, function(x){
  
  # grab the reference (Tr1) average composition and the focal composition
  temp = avg_rel[, c(grep(gsub("Tr2|Tr3|Tr5", "Tr1", x), colnames(avg_rel)),
                     grep(x, colnames(avg_rel))) ]
  
  # remove the invader and renormalize relative abundances
  temp[(rownames(temp) %in% c("MEJ082","RMX3_1b","LMC-P-011_S130")),2] = 0
  temp[,2] = temp[,2] / sum(temp[,2])
  
  # summarize in a data frame
  data.frame(
    sample = x, # the context
    isolate = sapply(names(which(temp[,1] > 0)), function(y){taxonomy$name[grep(y,taxonomy$isolate)]}), # the isolate
    rank = -(temp[which(temp[,1] > 0),1]) %>% rank(), # the rank of the relative abundance among the set
    before = temp[which(temp[,1] > 0),1], # the relative abundance in the Tr1 context
    after = temp[which(temp[,1] > 0),2] # the average relative abundance in the focal context
  )
  
}) %>% do.call(rbind.data.frame,.)

# remove cases where an isolate was lost from the community
results = results[which(results$after > 0),]

# difference/ratio of relative abundance after and before invasion as a metric of change
results$diff = results$after - results$before
results$ratio = results$after/results$before

# log ratio - this is the same as the proportional change but a log transformation improves the residuals of the ANOVA
anova = aov(data = results[-which(log(results$ratio) > (mean(log(results$ratio)) + 2*sd(log(results$ratio))) | 
  log(results$ratio) < (mean(log(results$ratio)) - 2*sd(log(results$ratio)))),], formula = log(ratio) ~ as.factor(rank)) # remove outlier ratios outside of 2SD

  qqPlot(anova$residuals)
  summary(anova)
  TukeyHSD(anova)

# is the decrease in abundance of rank-1 isolates significantly different from zero?
qqPlot(results$diff[which(results$rank == 1)])
t.test(x = results$diff[which(results$rank == 1)])
t.test(x = results$diff[which(results$rank == 9)])

#tiff("Figure3b.tiff", units="in", width=5, height=5, res=300)
ggplot(data = results, aes(x = as.factor(rank), y = log10(ratio))) + geom_boxplot(aes(group = rank), outlier.shape = NA) + 
  geom_jitter(aes(group = rank), alpha = 0.5, width = 0.25) + ylim(c(-1.6, 1.5)) +
  xlab("rank abundance") + ylab(expression(log["10"]~(post-invasion~"/"~pre-invasion~abundance))) + theme(text = element_text(size = 14)) +
  geom_text(x = 1, y = 1.55, label = "a", size = 6) + geom_text(x = 2, y = 1.55, label = "b", size = 6) + 
  geom_text(x = 3, y = 1.55, label = "b", size = 6) + geom_text(x = 4, y = 1.55, label = "b", size = 6) + 
  geom_text(x = 5, y = 1.55, label = "b", size = 6) + geom_text(x = 6, y = 1.55, label = "b", size = 6) +
  geom_text(x = 7, y = 1.55, label = "b", size = 6) + geom_hline(yintercept = 0, linetype = 2) +
  geom_text(x = 8, y = 1.55, label = "ab", size = 6) + geom_hline(yintercept = 0, linetype = 2) +
  geom_text(x = 9, y = 1.55, label = "ab", size = 6) + geom_hline(yintercept = 0, linetype = 2)
#dev.off()


# Figure 3a - compositional shifts pre/post invasion
T0 = read.csv(file = "T0_invasion.csv", header = TRUE)

### plotting - for a given structure where an invader is present, compare that to the composition of the un-invaded version
metadata$long_comm = gsub("C","Community ", metadata$community)

x = "Tr2_C12I2" # our nice example community
plot_temp = relabund[, c(grep(gsub("Tr2|Tr3|Tr5", "Tr1", x), colnames(relabund)),
            grep(x, colnames(relabund))) ] %>% melt()
colnames(plot_temp) = c("isolate","sample","relabund")
plot_temp = plot_temp[-which(plot_temp$relabund == 0),]
plot_temp$name = sapply(plot_temp$isolate, function(x){taxonomy$name[grep(x, taxonomy$isolate)]})
plot_temp$comm = sapply(plot_temp$sample, function(x){metadata$long_comm[grep(x, metadata$name)]})
plot_temp$sample = paste(substr(plot_temp$sample, 1, 3), substr(plot_temp$sample, 
            nchar(as.character(plot_temp$sample)), nchar(as.character(plot_temp$sample))), sep = "")
plot_temp$sample = gsub("Tr1"," Uninvaded_",plot_temp$sample)
plot_temp$sample = gsub(substr(x,1,3), # dude, this is bananas
            paste(gsub(" Invasion","",avg_invaders$treatment[grep(x, avg_invaders$sample)]),"_", sep = ""), plot_temp$sample)

# color-blind friendly palette
temp_palette <- c("#56B4E9", "#009E73", "#F0E442", "#0072B2", "#D55E00", "#CC79A7")

#tiff(file = "Figure3a.tiff", units="in", width=6.75, height=5, res=300)
  ggplot(data = plot_temp, aes(x = sample, y = relabund, fill = name)) + geom_col() + xlab("") + ylab("relative abundance") +
           theme(legend.title = element_blank(), text = element_text(size = 15), axis.text.x = element_text(angle = -60, hjust = 0)) +
           theme(legend.position = "top") + ggtitle(expression(Invader~(italic("P. viridiflava"))~excluded)) +
           guides(fill = guide_legend(nrow = 2)) + scale_fill_manual(values=temp_palette)
#dev.off()  

rm(results, x, plot_temp, temp_palette)
################################################################################

# Compare the invasion treatments with the uninvaded controls in terms of community composition (beta-diversity)
################################################################################
# prune samples from treatments 1/2 that are very divergent in composition from the "average" sample
prunables = c("Tr1_C4I3b","Tr1_C9I3c","Tr1_C9I4a","Tr1_C9I4c","Tr1_C11I1a","Tr1_C11I1b","Tr1_C11I1c","Tr1_C11I2a","Tr1_C13I1a","Tr1_C13I1b","Tr1_C13I1c","Tr1_C13I2c",
              "Tr2_C4I3b","Tr2_C9I3c","Tr2_C9I4a","Tr2_C9I4c","Tr2_C11I1a","Tr2_C11I1b","Tr2_C11I1c","Tr2_C11I2a","Tr2_C13I1a","Tr2_C13I1b","Tr2_C13I1c","Tr2_C13I2c")

### pairwise PERMANOVA - is there a significant effect of invasion treatment on community composition relative to the uninvaded control
  # Late Invasion
  # make a pruned temporary version of relabund and metadata
  temp_relabund = relabund[,!(metadata$name %in% prunables)]
  temp_metadata = metadata[!(metadata$name %in% prunables),]
  # remove the invaders (to compare only the endemic community fraction)
  temp_relabund = temp_relabund[!(rownames(temp_relabund) %in% rownames(taxonomy)[which(taxonomy$invader != 0)]),] 
  temp_relabund = apply(temp_relabund, 2, function(x){x / sum(x)}) # reset relative abundances
  # just a single treatment at a time
  temp_metadata = temp_metadata[grep("Tr1|Tr2", colnames(temp_relabund)),]
  temp_relabund = temp_relabund[,grep("Tr1|Tr2", colnames(temp_relabund))]
  # PERMANOVA
  temp = temp_relabund %>% t() %>% vegdist(., index = "bray")
  adonis2(data = temp_metadata, formula = temp ~ treatment, permutations = 999, strata = temp_metadata$community)
  
  # Initial Invasion
  # make a pruned temporary version of relabund and metadata
  temp_relabund = relabund[,!(metadata$name %in% prunables)]
  temp_metadata = metadata[!(metadata$name %in% prunables),]
  # remove the invaders (to compare only the endemic community fraction)
  temp_relabund = temp_relabund[!(rownames(temp_relabund) %in% rownames(taxonomy)[which(taxonomy$invader != 0)]),] 
  temp_relabund = apply(temp_relabund, 2, function(x){x / sum(x)}) # reset relative abundances
  # just a single treatment at a time
  temp_metadata = temp_metadata[grep("Tr1|Tr3", colnames(temp_relabund)),]
  temp_relabund = temp_relabund[,grep("Tr1|Tr3", colnames(temp_relabund))]
  # PERMANOVA
  temp = temp_relabund %>% t() %>% vegdist(., index = "bray")
  adonis2(data = temp_metadata, formula = temp ~ treatment, permutations = 999, strata = temp_metadata$community)
  
  # Early Invasion
  # make a pruned temporary version of relabund and metadata
  temp_relabund = relabund[,!(metadata$name %in% prunables)]
  temp_metadata = metadata[!(metadata$name %in% prunables),]
  # remove the invaders (to compare only the endemic community fraction)
  temp_relabund = temp_relabund[!(rownames(temp_relabund) %in% rownames(taxonomy)[which(taxonomy$invader != 0)]),] 
  temp_relabund = apply(temp_relabund, 2, function(x){x / sum(x)}) # reset relative abundances
  # just a single treatment at a time
  temp_metadata = temp_metadata[grep("Tr1|Tr5", colnames(temp_relabund)),]
  temp_relabund = temp_relabund[,grep("Tr1|Tr5", colnames(temp_relabund))]
  # PERMANOVA
  temp = temp_relabund %>% t() %>% vegdist(., index = "bray")
  adonis2(data = temp_metadata, formula = temp ~ treatment, permutations = 999, strata = temp_metadata$community)


### Beta-diversity relative to uninvaded controls
# make a pruned temporary version of relabund and metadata ***
temp_relabund = relabund[,!(metadata$name %in% prunables)]
temp_metadata = metadata[!(metadata$name %in% prunables),]
# remove the invaders (to compare only the endemic community fraction) ***
temp_relabund = temp_relabund[!(rownames(temp_relabund) %in% rownames(taxonomy)[which(taxonomy$invader != 0)]),] 
temp_relabund = apply(temp_relabund, 2, function(x){x / sum(x)}) # reset relative abundances
  
# gather the data appropriately
index = paste(metadata$community, metadata$invader, sep = "") %>% unique()
test = lapply(index, function(x){
  
  # pull out the Treatment 1 data as the references
  tr1 = temp_relabund[,grep(substr(x,1,nchar(x)-2), temp_metadata$sample[temp_metadata$treatment == "Tr1"])]
  
  # for each treatment, pull out the data from the relevant community/invader and calculate bray-curtis dissimilarity
  lapply(c("Tr2","Tr3","Tr5"), function(y){
    
    # pull out the relevant data for a given treatment
    # issue here - a given treatment is not always present, but it doesn't cause the code to crash
    if(length(which(temp_metadata$treatment[grep(x, temp_metadata$sample)] == y)) != 0){
      trx = temp_relabund[,grep(x, temp_metadata$sample)[which(temp_metadata$treatment[grep(x, temp_metadata$sample)] == y)]] %>% 
        as.data.frame()
      
      # calculate bray-curtis dissimilarity
      temp = cbind.data.frame(trx, tr1) %>% t() %>% vegdist(., index = "bray") %>% as.matrix()
      
      # remove the excess data
      temp = temp[(ncol(trx)+1):nrow(temp), 1:ncol(trx)]
      
      # average richness
      #rich = apply(trx, 2, function(z){length(which(z > 0))}) %>% mean(., na.rm = TRUE)
      
      # average growth on spent medium
      #growth = if(sum(spent$name %in% colnames(trx)) > 0){
      #  spent$growth[spent$name %in% colnames(trx)] %>% mean()
      #} else {NA}
      
      # format that information nicely
      #data.frame(comm_inv = x, treatment = y, avg = mean(temp), sd = sd(temp), richness = rich, growth = growth)
      
      ### testing ### do this for each column of temp instead of the average? increases the power
      # average richness
      rich = apply(tr1, 2, function(z){length(which(z > 0))}) %>% mean(., na.rm = TRUE)
      
      # average growth on spent medium
      growth = sapply(colnames(trx), function(z){ if(z %in% spent$name){spent$growth[spent$name %in% z]} else {NA} })
      
      # format that information nicely
      data.frame(comm_inv = x, treatment = y, avg = colMeans(as.data.frame(temp)), sd = apply(as.data.frame(temp), 2, sd), 
                 richness = rich, growth = growth)
      ###
      
    }
    
  }) %>% do.call(rbind.data.frame, .)
  
}) %>% do.call(rbind.data.frame, .)
test$comm = gsub("I1|I2|I4", "", test$comm_inv) %>% gsub("C","Community_",.)
test$invader = substr(test$comm_inv, nchar(test$comm_inv)-1, nchar(test$comm_inv))
test$treat = sapply(test$treatment, function(x){ if(x == "Tr2"){"Late"} else { if(x == "Tr3"){"Initial"} else {"Early"} } })

# bring in richness of the relevant invaded community
test$name = rownames(test)
test = merge(test, rich, by = "name")

# ANOVA comparing average dissimilarity between each invasion treatment relative to the uninvaded control communities
# Supplementary Table 3
test %>% group_by(treatment) %>% summarise(avg = mean(avg))
anova = aov(data = test, formula = avg ~ treat)
  summary(anova)
  leveneTest(data = test, avg ~ treat) # confirm homogeneity of variance
  qqPlot(anova)
  TukeyHSD(anova)

# Supplementary Table 4
# ANCOVAs considering growth on spent media and pre-invasion community richness
anova = aov(data = test, formula = avg ~ treat + growth)
  summary(anova)
  qqPlot(anova)
anova = aov(data = test, formula = avg ~ treat + richness.y)  
  summary(anova)
  qqPlot(anova)

rm(temp,test,anova,index,temp_relabund,temp_metadata,prunables)
################################################################################

# how often did a successful invasion result in the loss of a community member?
################################################################################
### i.e., when an invader was successful, how often did it displace a community member versus just slotting in?
  # only look at this for the no-invader (T7) control and late invasion (T7) because that is a direct comparison of the same communities before and after invasion

# isolate the data by treatment
tr1 = relabund[,which(metadata$treatment == "Tr1")] # no invasion
tr2 = relabund[,which(metadata$treatment == "Tr2")] # late invasion (of the uninvaded samples)

# I need to line them up 1:1 btw treatments 1 & 2 and 4 & 5
tr1 = tr1[,gsub("Tr1_","",colnames(tr1)) %in% gsub("Tr2_","",colnames(tr2))]
tr2 = tr2[,gsub("Tr2_","",colnames(tr2)) %in% gsub("Tr1_","",colnames(tr1))]
  all(gsub("Tr2_","",colnames(tr2)) == gsub("Tr1_","",colnames(tr1))) # sanity check

# make them presence/absence
tr1[which(tr1 > 0)] = 1
tr2[which(tr2 > 0)] = 1

# for each sample, how many isolates were detected in the uninvaded samples but not in the invaded samples?
exclude = data.frame(
  sample = colnames(tr1),
  n_excl = sapply(1:ncol(tr1), function(x){ (!(rownames(tr1)[which(tr1[,x] == 1)] %in% rownames(tr2)[which(tr2[,x] == 1)])) %>% sum() }),
  n_change = sapply(1:ncol(tr1), function(x){ length(which(tr2[,x] == 1)) - length(which(tr1[,x] == 1)) }),
  inv_present = sapply(1:ncol(tr1), function(x){ rownames(tr2)[which(tr2[,x] == 1)] %in% 
      c("MEJ082","RMX3_1b","LMC-P-011_S130") %>% sum() }) )

# when an invader was present, how often did it result in exclusion of a resident community member?
length(which(exclude$n_excl[which(exclude$inv_present == 1)] > 0)) / length(which(exclude$inv_present == 1))
################################################################################
```
